# Conformational resolution of nucleotide cycling and effector interactions for multiple small GTPases in parallel

**DOI:** 10.1101/577437

**Authors:** Ryan C. Killoran, Matthew J. Smith

## Abstract

Small GTPase proteins alternatively bind GDP/GTP guanine nucleotides to gate signaling pathways that direct most cellular processes. Numerous GTPases are implicated in oncogenesis, particularly three RAS isoforms HRAS, KRAS and NRAS, and the RHO family GTPase RAC1. Signaling networks comprising small GTPases are highly connected, and there is evidence of direct biochemical crosstalk between the functional G-domains of these proteins. The activation potential of a given GTPase is contingent on a co-dependent interaction with nucleotide and a Mg^2+^ ion, which bind to individual variants via distinct affinities coordinated by residues in the nucleotide binding pocket. Here, we utilize a selective-labelling strategy coupled with real-time nuclear magnetic resonance (NMR) spectroscopy to monitor nucleotide exchange, GTP hydrolysis and effector interactions of multiple small GTPases in a single complex system. We provide new insight on nucleotide preference and the role of Mg^2+^ in activating both wild-type and oncogenic mutant enzymes. Multiplexing reveals GEF, GAP and effector binding specificity in mixtures of GTPases and establishes the complete biochemical equivalence of the three related RAS isoforms. This work establishes that direct quantitation of the nucleotide-bound conformation is required to accurately resolve GTPase activation potential, as GTPases such as RALA or the G12C mutant of KRAS demonstrate fast exchange kinetics but have a high affinity for GDP. Further, we propose that the G-domains of small GTPases behave autonomously in solution and nucleotide cycling proceeds independent of protein concentration but is highly impacted by Mg^2+^ abundance.

## Introduction

Small GTPases are a class of critical hub proteins, responsible for controlling both the direction and intensity of cell signals by acting as ‘molecular switches’ (1–3). The RAS and RHO subfamilies constitute about one third of all RAS superfamily GTPases (4), and are key regulators of both normal and oncogenic cellular processes including proliferation and migration. Archetypally, small GTPases interconvert between active GTP-bound and inactive GDP-bound states. Nucleotide exchange and GTP hydrolysis occur intrinsically, however, exchange and hydrolysis rates are catalyzed by guanine nucleotide exchange factors (GEFs) or GTPase activating proteins (GAPs), respectively. Contributions from intrinsic versus catalyzed exchange *in vivo* are not well understood. In the active state, GTPases bind directly to downstream effector proteins *via* specialized recognition domains, predominantly <u>R</u>AS-<u>b</u>inding <u>d</u>omains (RBDs) for RAS GTPases and CRIB motifs for RHO GTPases (5, 6).

Within this enzyme superfamily, the most heavily studied are three highly related RAS isoforms: HRAS, KRAS and NRAS. These small GTPases are critical mediators of signaling networks that stimulate cell growth and proliferation, including the MAPK and PI3K/mTOR pathways. Oncogenic mutations at codons 12, 13 and 61 of the *HRAS, KRAS* and *NRAS* genes are amongst the most frequent genetic mutations in human cancers (7). The three RAS proteins share 80% sequence identity and are co-expressed in most cell types. Structurally, they share a nearly identical tertiary fold. As such, these proteins were initially considered functionally redundant, yet multiple lines of evidence support functional specificity: *RAS* genes exhibit different transforming potential (8–10), are distinctly mutated in cancers (11, 12), exhibit unique sensitivities to GEFs (13) and are localized to discrete subcellular locales (14). Whether genuine biochemical variations in the core G-domains of these proteins contribute to these observed biological differences remains an open question.

Existing approaches to measure small GTPase kinetics include time course-HPLC, release of ^32^P_i_ from GTP-^32^P_i_ and, most commonly, release or uptake of fluorescent nucleotide analogs. These are crucial assays used to decipher how much activated GTPase subsists *in vivo*. Unfortunately, several important drawbacks exist with these methods, foremost that fluorescently-tagged analogs can impact reaction kinetics (15). Indeed, the indirect nature of these approaches are a shortcoming that has led to improper conclusions concerning the rate of wild-type and oncogenic RAS mutant nucleotide exchange and its overall on/off state (16, 17). To accurately quantify the activation state of GTPases requires consideration of relative nucleotide affinity (i.e. preference of a given GTPase to bind GDP or GTP), the availability of Mg^2+^ cofactor, and the potential impact of multimer formation or membrane interactions. This would take into consideration the growing evidence that RAS GTPases dimerize (18, 19), which would be intensified at high protein concentrations such as those in membrane nanoclusters.

Recently, real-time nuclear magnetic resonance (RT-NMR) experiments have been adapted to quantitate small GTPase activity (20, 21). As GTPases undergo major conformational change upon binding to GDP or GTP, successive collection of ^1^H-^15^N HSQC spectra allows for kinetic analyses of exchange or hydrolysis. Importantly, these assays do not require fluorescent nucleotide analogs nor any chemical modification of the GTPase. Further, as NMR assays are functional over a wide range of protein concentrations and even on membrane-tethered GTPase (22), they can be used to probe the functional impact of proposed RAS GTPase oligomerization. To strengthen the RT-NMR approach, it is now possible to multiplex these assays (23), allowing quantification of activation states for several GTPases monitored simultaneously in real-time.

We employ here a multiplexed RT-NMR approach to study the full GTPase nucleotide exchange and hydrolysis cycle, as well as specificity of effector RBD binding. Using a selective-labelling strategy we employ RT-NMR to concurrently measure kinetics and effector specificity of the three related RAS isoforms, across RAS and RHO subfamily members, and between cancer-associated mutations of RAS and RAC1 and wild-type counterparts. The data improve our understanding of the complexity and inter-connectedness of small GTPases in the context of cell signaling, particularly the impact of relative nucleotide affinity, Mg^2+^ availability and cross-talk mechanisms.

## Results

### Selective Isotopic GTPase Labelling and Intrinsic Nucleotide Cycling

To juxtapose structure-function data for multiple small GTPases in parallel, we selected six enzymes from the RAS and RHO subfamilies (**Fig. 1A**). Structures of these proteins demonstrate the exceptional similarity of their tertiary folds (**Fig. 1B**). Identical sizes, shapes and enzymatic functions makes *in vitro* profiling of numerous GTPase activities together a massive challenge. For NMR-based analyses, uniformly labeled samples of multiple GTPases results in excessive crowding of resonance peaks, as demonstrated in **Fig. 1C** using HRAS, KRAS and NRAS. To observe multiple GTPases concurrently required a selective isotopic labelling approach with single ^15^N amino acids. **Fig. 1D** and **1E** show isotopically labeled RAS isoforms (^15^N-Ile HRAS, ^15^N-Thr KRAS and ^15^N-Leu NRAS) in both the GDP-bound and GTP-bound conformation (using non-hydrolyzable GMPPNP). The reduction in spectral complexity allowed us to monitor nucleotide exchange, GTP hydrolysis or effector binding in mixtures of even the most highly related GTPases.

**Fig. 1.**
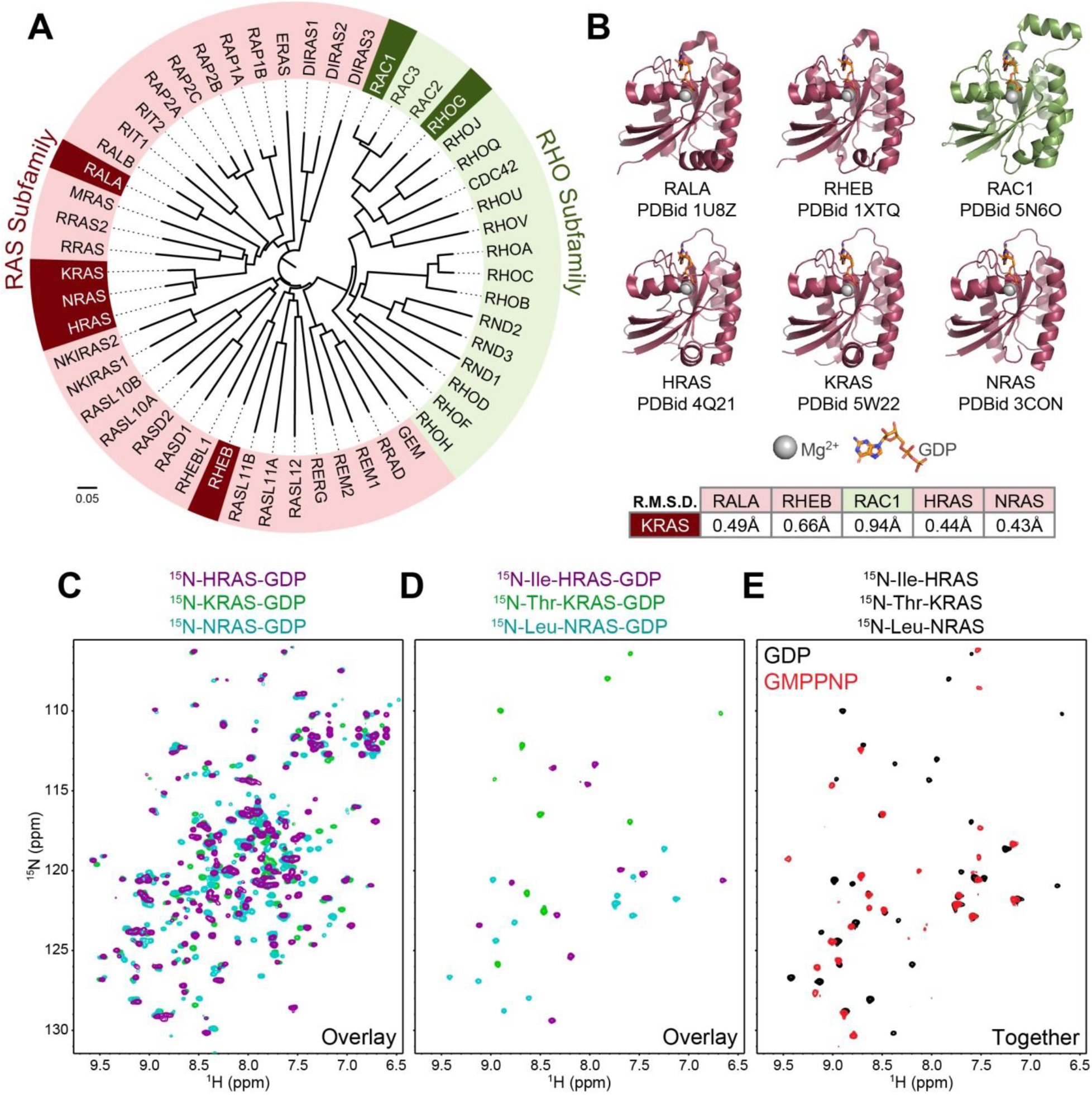
NMR spectroscopy to monitor activity of multiple, highly similar small GTPases simultaneously. **A)** Amino acid sequence alignment of the complete RAS and RHO subfamily small GTPases. GTPases used in this study are highlighted. **B)** Available structures of GTPases used in this study. R.M.S.Ds between these GTPases and KRAS demonstrate structural homology. **C)** HSQC overlay of uniformly ^15^N labeled HRAS, KRAS and NRAS loaded with GDP. **D)** HSQC overlay of selectively labeled RAS isoforms loaded with GDP: ^15^N Ile-labeled HRAS (purple), ^15^N-Thr labeled KRAS (green) and ^15^N-Leu labeled NRAS (blue). **E)** Mixture of selectively labeled RAS isoforms from *D)*, loaded with GDP (black spectrum) and GMPPNP (red spectrum).

We performed exchange and hydrolysis assays on individual selectively-labeled GTPases to profile their kinetic activity in isolation. NMR spectra and nucleotide exchange plots for ^15^N-Tyr HRAS, ^15^N-Thr KRAS, ^15^N-Leu NRAS, ^15^N-Leu RALA, ^15^N-Val RHEB, ^15^N-Val RHOG and ^15^N-Leu RAC1 are displayed in **Supplemental Fig. S1** and **S2A-G**. Kinetics for these reactions are detailed in **Supplemental Table S1** (for all individually measured GTPase kinetics). Exchange was initiated by the addition of a 10:1 molar ratio of the GTP analog GTPγS, used to reflect the cellular GTP/GDP ratio. Measured exchange rates for HRAS match what has been calculated by RT-NMR using uniformly labeled samples (21). As individual GTPases reach an activation plateau (i.e. nucleotide exchange no longer proceeds), they exhibit differential GTPγS-loading that reflects their activation state at equilibrium. We report these data in **Supplemental Table S1** as “% Activated”. Further, as we control the GTPγS:GDP ratio at 10:1 we can also calculate the preference of these GTPases to each nucleotide, reported as “GDP Preference”. We observed the three RAS isoforms and RHEB exhibited a much higher ratio of GTPγS-bound (∼65-75%) in these conditions compared to RALA (38%) or RHOG (49%), a direct reflection of differential nucleotide affinities. We next measured intrinsic GTP hydrolysis for each RAS isoform (^15^N-Tyr HRAS, ^15^N-Thr KRAS and ^15^N-Leu NRAS) (**Supplemental Fig. S2H-J**). The measured rate for HRAS matches what was previously determined on uniformly labeled protein (21), and NRAS and KRAS exhibited near identical rates. Overall, the selective-labeling approach provides an opportunity to precisely measure GTPase activity while reducing spectral complexity.

### Dependence of Nucleotide Exchange on Mg^2+^

To begin assay of multiplexed GTPases, we first focused on the three highly related RAS isoforms. There has been little attempt to study differences in nucleotide exchange and/or GTP hydrolysis rates for the RAS isoforms in their individual contexts, however, one study found that the intrinsic rates of GTP catalysis differed across isoforms using ^32^Pi-GTP single turnover (24). Due to the inefficiency of selective-labeling, monitoring HSQC chemical shifts of the three GTPases required each at a concentration of 300 μM. Interestingly, this meant we could assay GTPase activity at concentrations that should drive RAS dimerization. The existence of KRAS homodimers remains contentious, with proposed dissociation binding constants ranging from low µM (25) to mM (26). Recent biophysical data dispute the existence of RAS dimers even at high concentrations (27, 28). Performing these assays at total GTPase concentrations approaching 1 mM would resolve whether G-domain oligomers can influence RAS activation in solution. Initial multiplexed nucleotide exchange assays (GDP-to-GTPγS) at a 10:1 molar excess of GTPγS showed exchange rates substantially faster than those measured for GTPases individually at comparatively lower concentrations. To resolve this, we considered that exchange assays with each RAS isoform alone at various concentrations (150, 250 or 350 μM), showed rates increase with protein concentration (**Fig. 2A-C**). Mg^2+^ cofactor is absolutely required for nucleotide binding to RAS GTPases, and both RAS (29) and RHO (30) nucleotide exchange is highly dependent on Mg^2+^ concentration. The majority of GTPase kinetic studies use MgCl_2_ at a steady concentration of 5 mM, and we postulated that increasing protein concentrations lower the [Mg^2+^]:[GTPase] ratio, leading to Mg^2+^ scarcity and faster exchange rates. To test this, we measured nucleotide exchange of NRAS at 200 µM or 350 µM in 5 mM MgCl_2_, again showing that the rate increases at a higher concentration (**Fig. 2D**). When we repeated the assay using 15 mM MgCl_2_, exchange rates were identical. Thus, Mg^2+^ availability is a key determinant of GTPase exchange, which otherwise proceeds independent of protein concentration even approaching 1 mM.

**Fig. 2.**
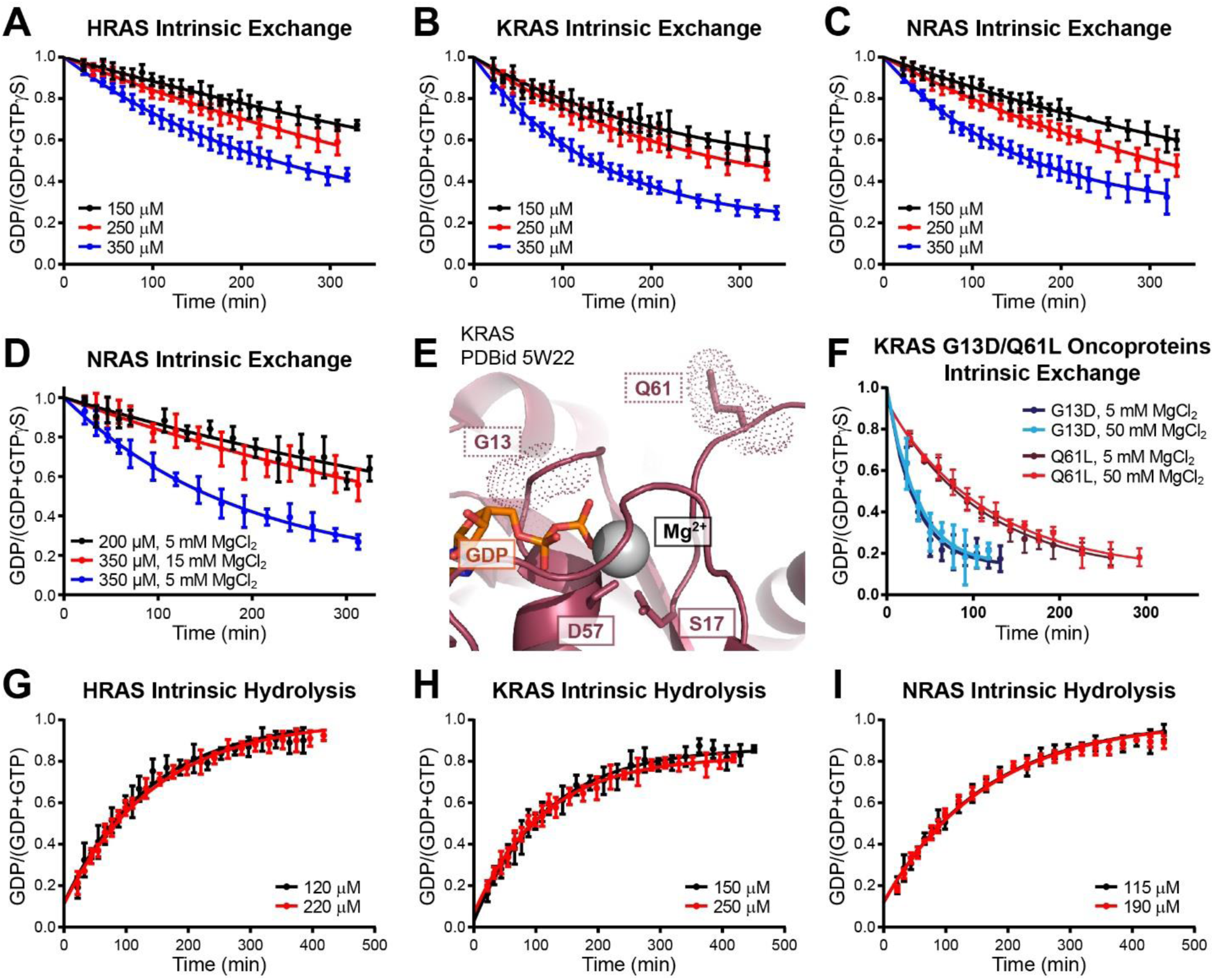
Mg^2+^ dependence of nucleotide exchange. **A-C)** Intrinsic nucleotide exchange assays performed on *A)* HRAS, *B)* KRAS and *C)* NRAS at increasing concentrations of uniformly labeled GTPase at a 10:1 GTPγS:GTPase molar ratio. **D)** Intrinsic nucleotide exchange assays with NRAS at two concentrations (200 and 350 μM). Assays performed at 350 μM were performed in 5 mM and 15 mM MgCl_2_. **E)** Co-ordination of Mg^2^+ in GDP-loaded KRAS. Positions of mutation hotspots G13 and Q61 are highlighted. **F)** Intrinsic nucleotide exchange assays performed on KRAS G13D and Q61L at a 10:1 GTPγS. **G-I)** Intrinsic GTP hydrolysis assays of *G)* HRAS, *H)* KRAS and *I)* NRAS performed at varying concentrations of uniformly labeled GTPase.

Interestingly, there are several recurrent oncogenic mutants of RAS that have been determined to function by *via* rapid intrinsic exchange (21). As these mutations lie proximal to the nucleotide/Mg^2+^ binding pocket we were curious if increasing [Mg^2+^] may slow their intrinsic exchange rate. We purified isotopically labeled KRAS proteins of two fast exchange mutants, G13D (found within the P-loop) and Q61L (in the switch II region) (21, 31) (**Fig. 2E**). We performed nucleotide exchange assays at either 5 or 50 mM MgCl_2_ (**Fig. 2F**). There was no Mg^2^+ dependence on the exchange rate of either mutant, indicating these amino acid mutations likely alter nucleotide affinity rather than disrupt coordination of the Mg^2+^ ion.

Finally, we tested whether increasing concentrations of RAS with a steady concentration of Mg^2+^ would lead to differences in GTP hydrolysis. Neither KRAS, HRAS nor NRAS exhibited altered GTP hydrolysis rates with increasing protein concentration at a constant [Mg^2+^] of 5 mM (**Fig. 2G-I**). This is consistent with the absence of competing nucleotide in these assays, and with data suggesting GDP-binding is more dependent on Mg^2+^ than is GTP binding (32).

### Multiplexed Nucleotide Exchange Assays

With a strategy for selective amino acid labelling and conditions optimized for simultaneously monitoring multiple GTPase activities, we performed a series of multiplexed nucleotide exchange assays. **Fig. 3A-C** shows HSQC spectra depicting the amino acid labelling strategy and consequent GDP/GTPγS-bound peaks used to concurrently measure exchange rates for HRAS, KRAS and NRAS. Temporal resolution of chemical shifts and plots for intrinsic nucleotide exchange of these RAS isoforms are in **Fig. 3D/E**. For this experiment, we utilized a [GTPγS]:[GTPase] of 8:1 and 15 mM MgCl_2_. Each isoform was at a concentration of 300 μM. Mixing multiple GTPases at these concentrations did not lead to obvious binding or higher order complexes, as no significant chemical shift perturbations or peak broadening was observed. Intrinsic exchange rates for each of the three isoforms were not changed from those measured for HRAS, KRAS and NRAS alone at lower concentrations. Kinetic analyses and parameters for all multiplexed assays are detailed in **Supplemental Table S2**. All three GTPases were 80-90% activated at equilibrium and demonstrate a minor preference for GDP over GTPγS. We can conclude that neither the presence of alternative RAS isoforms nor high concentrations of these GTPases significantly impact their activation state.

**Fig. 3.**
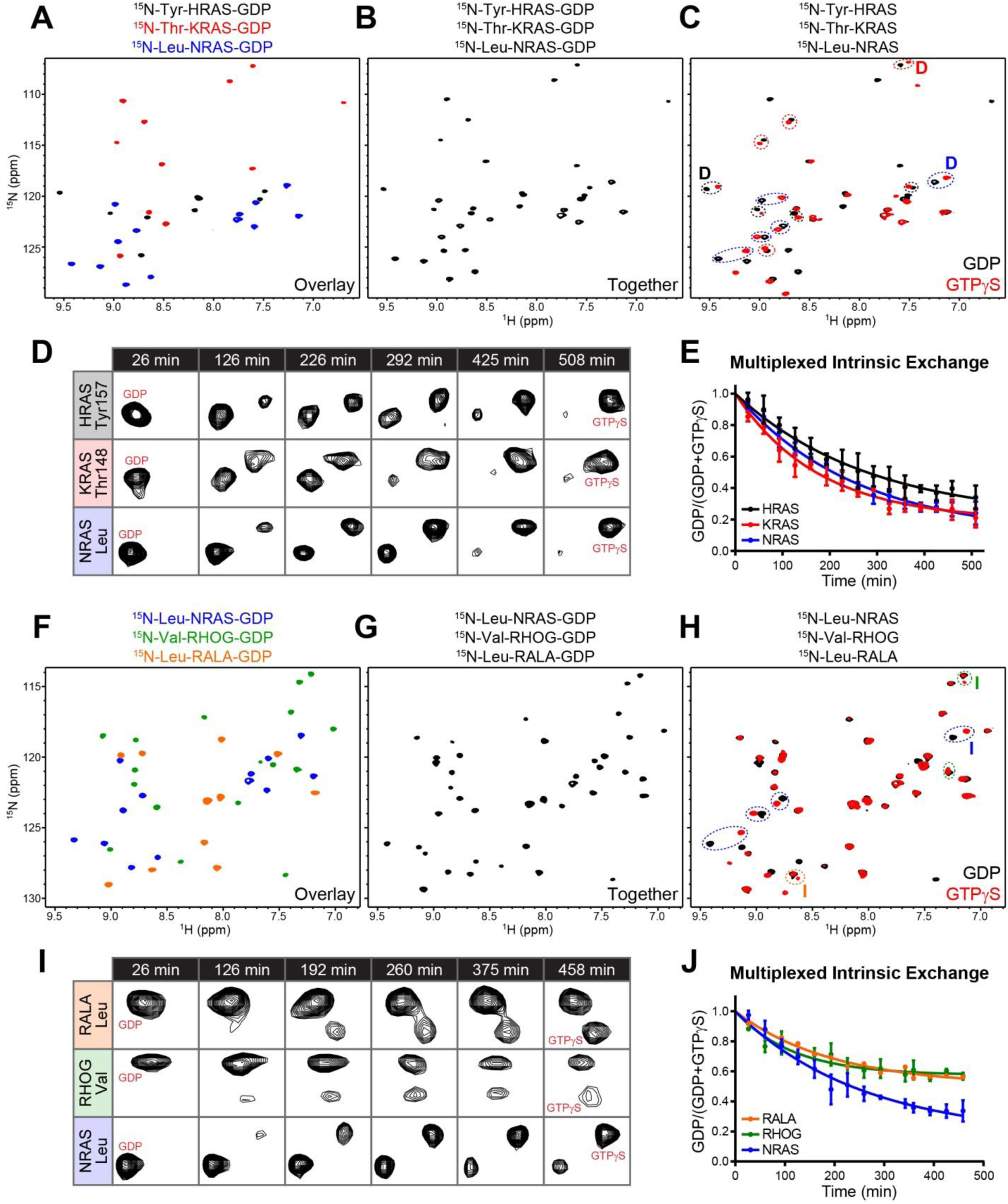
Multiplexed nucleotide exchange assays with small GTPases **A-C)** Multiplexed assay for the three RAS isoforms. *A)* Overlay of GDP-loaded ^15^N-Tyr HRAS, ^15^N-Thr KRAS and ^15^N-Leu NRAS collected at 300 μM. *B)* Multiplexed mixture of GDP-loaded ^15^N-Tyr HRAS, ^15^N-Thr KRAS and ^15^N-Leu NRAS each at 300 μM. *C)* Multiplexed nucleotide exchange of specifically labeled H-, K- and NRAS (black spectrum=GDP, red spectrum=GTPγS). Several peaks exhibiting chemical shift perturbation are circled (black=HRAS, red=KRAS, blue=NRAS). **D)** Representative resonances (labeled ‘D’ in panel *C*) for HRAS, KRAS and NRAS across multiple time points of multiplexed exchange. **E)** Intrinsic exchange curves of multiplexed HRAS, KRAS and NRAS initiated by the addition of 8:1 molar ratio of GTPγS in 15 mM MgCl_2_. **F-H)** Multiplexed scheme for monitoring exchange of NRAS, RHOG and RALA. *F)* Overlay of GDP-loaded ^15^N-Leu NRAS, ^15^N-Val RHOG and ^15^N-Leu RALA at 300 μM. *G)* Multiplexed mixture of ^15^N-Leu NRAS, ^15^N-Val RHOG and ^15^N-Leu RALA each at 300 μM. *H)* Multiplexed exchange of specifically labeled NRAS, RHOG and RALA (black=GDP, red=GTPγS). Several peaks exhibiting perturbation are circled (blue=NRAS, green=RHOG, orange=RALA). **I)** Representative resonances for NRAS, RHOG and RALA (labeled ‘I’ in panel *H*) across multiple time points of the multiplexed exchange. **J)** Intrinsic exchange curves of multiplexed NRAS, RHOG and RALA initiated by the addition of 8:1 GTPγS in 15 mM MgCl_2_.

We next examined the NRAS GTPase in parallel with a related GTPase, RALA, and a RHO subfamily GTPase, RHOG. **Fig. 3F-J** depicts the labelling strategy and multiplexed exchange. RALA and RHOG exhibited slightly faster exchange rates than NRAS (1.4-fold and 1.3-fold, respectively), but their % activation at equilibrium is significantly lower than that of NRAS (84% GTPγS-bound for NRAS, 46% RALA, and 49% RHOG). This reveals that RALA and RHOG have a greater preference for GDP over GTPγS compared to NRAS. These data highlight the need to directly monitor the nucleotide-bound state when quantifying GTPase activation, as kinetic rates alone can significantly misrepresent the total activated GTPase in a given system.

### Multiplexed GEF and GAP Assays

Our multiplexed NMR strategy presents an opportunity to observe the effects and specificity of exchange-promoting GEFs and hydrolysis-activating GAPs on mixtures of small GTPases (**Fig. 4A**). We performed a GEF assay in a mixture of the three isoforms of RAS using a catalytic domain from the major RAS regulator SOS1 (SOS_cat_ (21, 33)), added at a molar ratio of 1:15000 (versus [total GTPase]) (**Fig. 4B**). We calculated similar increases in the SOS-catalyzed exchange rate for all three isoforms, (4.5-fold increase for HRAS, 3.3-fold for KRAS and 3-fold for NRAS) indicating that they have comparable sensitivities to this GEF.

**Fig. 4.**
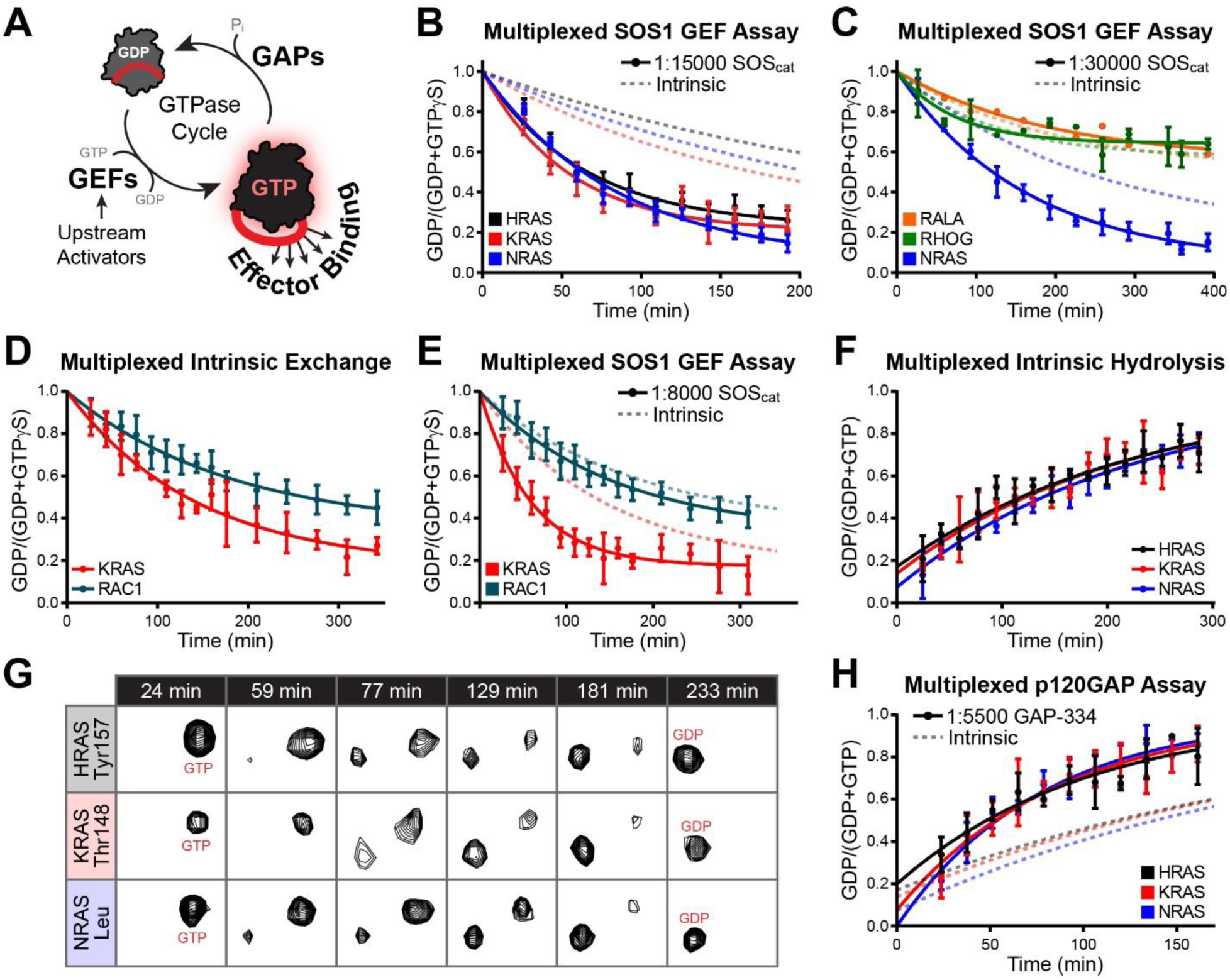
Multiplexed GTPase assays to profile GEFs and GAPs. **A)** The GTPase cycle; nucleotide exchange is enhanced by GEFs and GTP hydrolysis by GAPs. **B)** Multiplexed exchange curves of HRAS, KRAS and NRAS with recombinant SOS_cat_. Exchange was initiated by 8:1 GTPγS in 15 mM MgCl_2_. **C)** Nucleotide exchange curves of multiplexed NRAS, RALA and RHOG with recombinant SOS_cat_ under the same conditions. **D)** Intrinsic exchange of multiplexed KRAS and RAC1 at 300 μM each initiated by 10:1 molar ratio of GTPγS in 10 mM MgCl_2_. **E)** Multiplexed exchange of KRAS and RAC1 with recombinant SOS_cat_ demonstrates the GEF specificity. **F)** Multiplexed GTP hydrolysis of HRAS, KRAS and NRAS at 125 μM each. **G)** Representative resonances for HRAS, KRAS and NRAS across multiple time points of a multiplexed intrinsic hydrolysis assay. **H)** GTP hydrolysis curves of multiplexed HRAS, KRAS and NRAS with GAP-334.

To observe SOS specificity, we performed the same assay using multiplexed NRAS, RALA and RHOG (**Fig. 4C**). In the presence of SOS_cat_, we observed a significant increase in the exchange rate of NRAS relative to intrinsic and no effect on the exchange rates of RALA or RHOG. We next performed a multiplexed assay using KRAS and RAC1, as SOS is reported to have GEF activity towards both. The activating GEF domains differ for each GTPase: the REM-CDC25 domains (SOS_cat_) mediate activity towards RAS (33, 34), and the DH-PH domains activate RAC1 (35, 36). The intrinsic exchange of KRAS proceeds 1.3-fold faster than that of RAC1, with % activated at equilibrium measured at 83% and 65%, respectively (**Fig. 4D**). Upon addition of SOS_cat_, KRAS nucleotide exchange increases 2.5-fold while RAC1 rates remained unchanged (**Fig. 4E**). Notably, the nucleotide bound ratio of KRAS at equilibrium is not altered by the presence of SOS (83%), supporting a model whereby GEFs do not actively exchange nucleotide but function by a passive mechanism.

We next employed our approach to measure GAP activation of GTPase hydrolysis by performing multiplexed assays with the three RAS isoforms. Calculated intrinsic hydrolysis rates for HRAS, KRAS and NRAS were indistinguishable when measured in tandem (**Fig. 4F/G**). In the presence of recombinant GAP-334 domain from the major RAS regulator p120GAP (added at a 1:5500), hydrolysis rates of each isoform were uniformly increased (2.3-fold HRAS, 2.6-fold KRAS and 2.9-fold NRAS (**Fig. 4H**)). These results indicate that the three isoforms share sensitivity to GAP activation, in addition to having comparable rates of intrinsic hydrolysis.

### Multiplexed GTPase Assays with Oncoproteins

There are conflicting reports on wild-type RAS isoforms influencing the transformation potential of oncogenic mutants, and little is known about cross-talk between wild-type and mutant GTPases from a biophysical standpoint. We used multiplexed RT-NMR to concurrently monitor nucleotide exchange and GTP hydrolysis of wild-type RAS isoforms and several oncogene-derived mutants (**Fig. 5A**: KRAS-G12C, KRAS-Q61L, and HRAS-G12V). HSQC spectral resolution of the activation state and nucleotide exchange assays on the individual oncoproteins are presented in **Supplemental Fig. S3A-C** and **S4A-C**. HRAS G12V and KRAS G12C have intrinsically slow nucleotide exchange compared to their wild-type counterparts, while KRAS Q61L exhibits rapid intrinsic exchange. This was also observed when these oncoproteins were multiplexed with the matched wild-type, whereby KRAS G12C intrinsically exchanges at a rate 2.9-fold slower than wild-type, HRAS G12V at a rate 1.6-fold slower, and KRAS Q61L 1.6-fold faster (**Supplemental Fig. S4D-F**). Significantly, the KRAS G12C variant reaches equilibrium at only 67% activated and HRAS G12V at only 36%. In contrast, KRAS Q61L reaches 87% activation under these conditions. We extended these analyses to measure GEF activity by adding SOS_cat_. The KRAS G12C mutant had a 1.5-fold slower exchange rate in the presence of SOS_cat_ than wild-type KRAS, similar to the intrinsic difference (**Fig. 5B**). A multiplexed GEF assay with wild-type KRAS and the Q61L variant again demonstrated that Q61L exchanges at a faster rate than wild-type (1.3-fold) and reaches equilibrium at nearly 90% GTPγS-bound (**Fig. 5C**). These data suggest that there is no biophysical interplay between wild-type and oncogenic RAS GTPases at even high concentrations of G-domain. They suggest that large pools of the RAS G12V or G12C variants likely remain GDP-bound even in the presence of activating GEF.

**Fig. 5.**
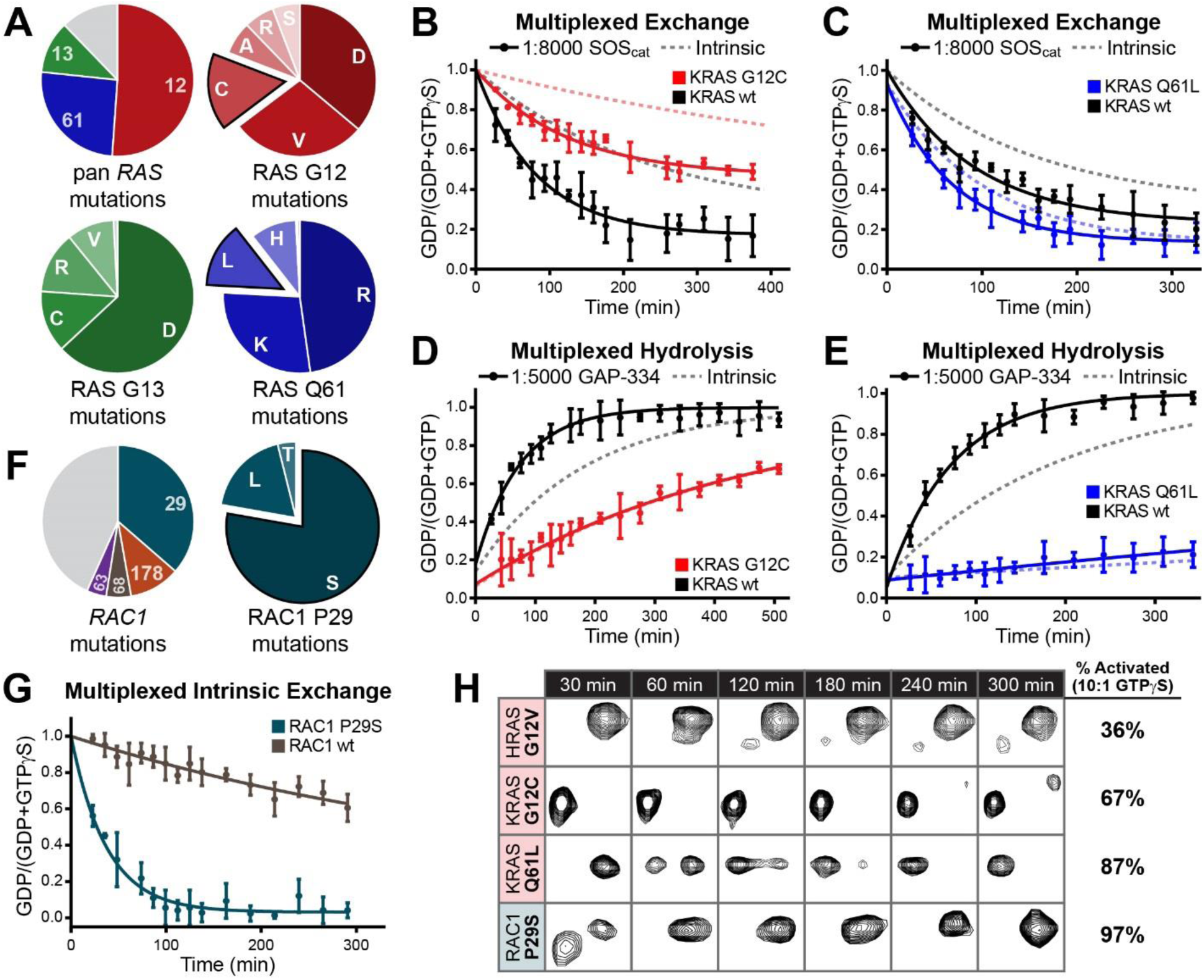
Multiplexed GTPase analyses of wild-type RAS or RAC1 variants complexed with oncogenic mutants. **A)** RAS mutations are recurrently found at three hotspots: codon 12, 13 and 61. Most common coding variants for each hotspot are displayed (complied from TCGA data). **B)** Multiplexed SOS_cat_ GEF assay using wild-type and G12C KRAS. 250 µM of each GTPase in 10 mM MgCl_2_,1:8000 SOS_cat_ and 10:1 GTPγS. **C)** Wild-type and Q61L KRAS GEF assay using the same conditions. **D)** Multiplexed wild-type and G12C KRAS GAP assay with GTP-loaded proteins and GAP-334 at 1:5000. **E)** Wild-type and Q61L KRAS GAP assay under the same conditions. **F)** Variant hotspots for RAC1 include codons 29, 63, 68 and 178. Most common coding variants for codon 29 are displayed (complied from TCGA data). **G)** Multiplexed intrinsic nucleotide exchange of wild-type and P29S RAC1 in 10 mM MgCl_2_ and 10:1 GTPγS. **H)** Snapshots of resonance transitioning from the GDP-bound state to GTPγS-bound state for HRAS G12V, KRAS G12C and Q61L and RAC1 P29S. Data are from multiplexed intrinsic exchange assays with the wild-type protein (RAS or RAC1).

Impaired GTP hydrolysis is a key biochemical defect in oncogenic RAS GTPases. We determined that KRAS G12C has a 2.6-fold slower intrinsic rate of hydrolysis than wild-type KRAS when measured individually (**Supplemental Fig. S4G**) or in a multiplexed assay (**Supplemental Fig. S4H**). The KRAS Q61L variant showed effectively no intrinsic GTP hydrolysis over a 10 hr time course (**Supplemental Fig. S4I**). To monitor rates in the presence of GAP, we added GAP-334 to multiplexed samples. The addition of GAP at 1:5000 did not affect the GTP hydrolysis rate of either G12C or Q61L KRAS (**Fig. 5D/E**), while the hydrolysis rate of wild-type KRAS increased 2.5-fold. Thus, the presence of neither wild-type KRAS nor GAP significantly alters the ability of these oncogenic mutants to hydrolyze GTP.

Genetic defects impacting small GTPase function in cancer are not limited to RAS proteins, so we sought to examine biophysical interplay between RAC1 and a RAC1 P29S mutant recurrently found in melanoma (**Fig. 5F**). A nucleotide exchange assay on the P29S variant alone demonstrated this oncoprotein intrinsically exchanges 7-fold faster than wild-type RAC1 (**Supplemental Fig. S4J**). A multiplexed approach with wild-type and P29S RAC1 provided the same result (**Fig. 5G**). Importantly, P29S RAC1 reaches almost 100% activation in these conditions, demonstrating a clear preference for binding GTPγS over GDP, while wild-type RAC1 shows a 6-fold preference for GDP. The utility of an approach that directly monitors GTPase conformation is fully demonstrated in **Fig. 5H** and **Supplemental Fig. S4K**. These NMR resonances demonstrate how poorly the HRAS G12V and KRAS G12C oncoproteins exchange over time in a 10-fold excess of GTPγS, as compared to their wild-type counterparts, KRAS Q61L or RAC1 P29S.

### Multiplexed Effector Binding Assays

The specificity of effector binding domains for small GTPases is a question of huge interest, and one that has not been well explored. Effectors targeting RAS subfamily GTPases typically comprise an RBD domain, and there are >50 potential RBDs in the human proteome (37). To directly observe effector specificity and competition for GTPase binding partners in a complex system we used NMR and multiple selectively labeled GTPases. We purified RBDs from two isoforms of the effector RAF kinases (ARAF and BRAF) and an RBD from the RAL effector RLIP76 (**Fig. 6A**). The RBD of ARAF was added to GMPPNP-loaded HRAS, KRAS and NRAS, and displayed uniform binding to all three isoforms as determined by peak broadening and chemical shift perturbations (**Fig. 6B**). Next, we titrated the RBD of BRAF and it also induced complete broadening across all three isoforms of RAS (**Fig. 6C**). To observe specificity across GTPase subfamilies, the BRAF RBD was titrated into a mixture of GMPPNP-loaded KRAS, NRAS and RHOG. Once again, severe peak broadening was observed for the KRAS and NRAS resonances, while peaks derived from RHOG were left unperturbed (**Fig. 6D**). The structurally unrelated RBD from RLIP76 was then titrated into a mixture of HRAS, KRAS and RALA. Only RALA peaks displayed peak broadening while HRAS and KRAS peaks were unperturbed (**Fig. 6E**). Finally, we looked to titrate an engineered monobody against RAS, NS1 (38), into a multiplexed mixture of HRAS, KRAS and NRAS. This monobody was designed to interact with the distal site (from the nucleotide-binding pocket) of HRAS and KRAS to prevent its dimerization/clustering on the membrane. Upon titration of NS1 into the three isoforms of RAS, peak broadening was observed only in HRAS and KRAS (**Fig. 6F**), while NRAS-specific chemical shifts were unperturbed. Overall, the multiplexed NMR approach is a powerful technique to observe interactions and binding specificities of multiple, unmodified proteins simultaneously.

**Fig. 6.**
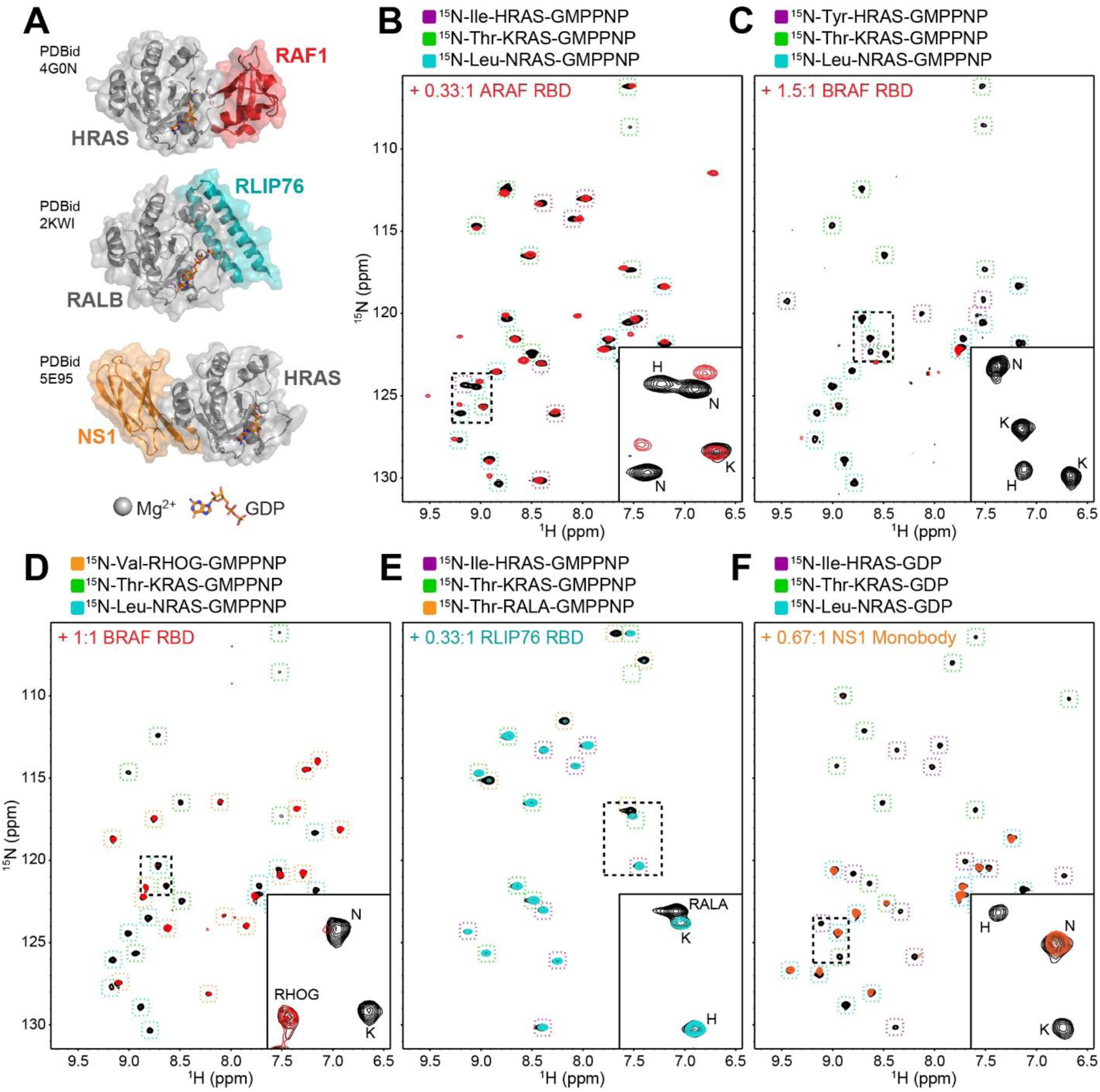
Multiplexed GTPase binding interactions. Peaks corresponding to each GTPase are highlighted by coloring described in legends (top). Displayed molar ratios are [effector]:[total protein]. **A)** GTPase/effector complex structures of HRAS and the RBD of RAF1, RALB and RLIP76 and the complex structure of HRAS with the engineered NS1 monobody. **B)** Selectively-labeled, GMPPNP-loaded H-, K- and NRAS (black spectrum) were mixed with ARAF RBD (red). **C)** Selectively-labeled, GMPPNP-loaded H-, K- and NRAS (black spectrum) were mixed with BRAF RBD (red). **D)** Selectively-labeled, GMPPNP-loaded KRAS, NRAS and RHOG (black spectrum) were mixed with BRAF RBD (red). **E)** Selectively-labeled, GMPPNP-loaded KRAS, HRAS and RALA (black spectrum) were mixed with RLIP76 RBD (blue). **F)** Selectively-labeled, GMPPNP-loaded H-, K- and NRAS (black spectrum) were mixed with NS1 monobody (orange).

## Discussion

Accurate data characterizing small GTPase nucleotide cycling are crucial to our understanding of both wild-type and mutant activation potential in cells. These are extremely challenging experiments that must consider co-dependent nucleotide and Mg^2+^ affinity, competing nucleotides and metal ions, and the distinct biochemical properties of a given GTPase. We have investigated the complete nucleotide cycle and effector interactions for multiple GTPases in parallel using a selective labeling approach coupled with RT-NMR. These multiplexed assays reveal details of GEF and GAP specificities, protein binding preferences, and nucleotide-dependent activation states that cannot be accurately assessed using conventional techniques.

Biochemical diversity in the three isoforms of RAS is an open question with large implications for developmental biology and cellular transformation. We find that intrinsic rates of exchange and hydrolysis are nearly identical across the three isoforms, as are their sensitivities to activation by SOS or inactivation by p120GAP. It appears that the core G-domains of HRAS, KRAS and NRAS are biochemically equivalent and that biological differences are likely determined by post-translational modification, subcellular localization or distinct effector/protein interactions. It will be interesting to determine if these results hold true for other RASGEF proteins such RASGRFs or RASGRPs (39, 40), and to examine GEF specificity against all 35 RAS subfamily GTPases (4).

Using high protein concentrations to profile GTPase kinetics revealed a strong dependence on Mg^2+^ for nucleotide exchange. The Mg^2+^ concentration in mammalian cells has been estimated at 17-20 mM, however, less than 5% of that is presumed free (41, 42). The cytoplasm (where membrane-tethered small GTPase proteins are exposed) is expected to have only 0.5-1 mM free Mg^2+^, significantly less than used in most *in vitro* kinetic assays. Several thoughtful experiments from nearly three decades ago estimated the Mg^2+^ affinity for wild-type HRAS at 2.8 µM (43), and resolved that high Mg^2+^ concentrations in RAS exchange assays significantly slow GDP dissociation rates. Moreover, there are intriguing data that Mg^2+^-GTP affinity for HRAS is higher than that of Mg^2+^-GDP (32). This would be consistent with our observations of increased nucleotide exchange in conditions of Mg^2+^ scarcity, but how this affects RAS activity in cells is unknown. We can speculate that high density RAS nanoclusters may be influenced by Mg^2+^ availability, which could act to promote nucleotide exchange. **Supplemental Table S3** complies the existing knowledge of RAS nucleotide binding and the little that is known about Mg^2+^ affinity. Most of these early data were generated by measuring retention of radiolabeled nucleotides on filters, but these detailed assessments of GTPase biochemistry and function should be reconsidered using modern approaches.

Multiple lines of evidence suggest functional interplay between wild-type and mutant RAS proteins, with claims that wild-type RAS can suppress oncogenic mutant activity (19, 44–46) and others that mutants promote wild-type RAS activation (47, 48). Multiplexing GTPase mutants with their wild-type counterpart G-domains provided an opportunity to explore direct biophysical effects on hydrolysis or exchange. Our results corroborate observations that mutations at G12 (Cys or Val) or Q61 (Leu) have unique deficiencies. The G12 oncoproteins exhibit very slow intrinsic exchange and hydrolysis rates, minor sensitivity to GEF and complete insensitivity to GAPs, while Q61L is a fast exchange mutant that exhibits virtually no hydrolysis of GTP. RAC1 P29S, the third most frequently observed hotspot mutation in melanoma (after NRAS and BRAF) (49, 50), is also a fast exchange mutant. We observed no significant cross-talk between wild-type and oncogenic mutant proteins, either in chemical shift perturbations upon incubation or by measuring nucleotide cycling kinetics.

Multiple selectively labeled proteins provide an excellent opportunity to directly measure binding competition in a complex *in vitro* system. Addition of the ARAF or BRAF RBDs to mixtures of the three RAS isoforms showed uniform binding to all three GTPases, while we could observe clear specificity differences using either distinct effector binding domains or distally related GTPases. While these NMR mixing experiments led to peak broadening in our system, improved approaches able to detect chemical shift perturbations will allow for calculation of affinities in a competitive setting. Future combination of multiplexed GTPases with multiple competing effectors (51) will facilitate study of ever-more complex *in vitro* systems that retain a capacity to deliver atomic-level, quantitative data for composite protein-protein interactions.

Precise and dynamic measurements of GTPase activation states are essential to elucidate their role in development and disease. Here, we have collectively demonstrated how selective isotopic labelling and RT-NMR offer improved measures of intrinsic, GEF and GAP enzymatic activities and effector binding specificities for multiple GTPases.

## Materials and Methods

### Plasmid Constructs

HRAS wild-type and G12V (residues 1-171), KRAS wild-type, G12C and Q61L (residues 1-171), the GAP-334 region of human p120GAP (residues 715-1047) and the SOScat domain of human SOS1 (residues 564-1049) are cloned into pET15b (Novagen / EMD Biosciences) with an N-terminal polyhistidine (His) tag. NRAS (residues 1-172), RHOG (residues 1-179), RALA (residues 1-183) and RAC1 wild-type and P29S (residues 1-177) were cloned into pDEST17 using Gateway Technology, with a thrombin cleavage site inserted between the His-tag and GTPase. Mouse RHEB (residues 1-169), human BRAF RBD (residues 150-233) and human RLIP76 RBD (residues 395-517) are cloned as a GST fusion in pGEX-2T. RHEB constructs were a kind gift from Dr. Vuk Stambolic (UHN Toronto). The NS1 monobody, expressed as a GST fusion protein, was gifted from Dr. Shohei Koide (NYU Langone).

### Protein Expression and Purification

For unlabeled GST-tagged proteins, proteins were expressed in *Escherichia coli* BL21 codon+ cells in LB media by induction with 0.25 mM isopropyl-b-D-thiogalactopyranoside at 16 °C overnight. Uniformly labelled ^15^N His or GST-tagged proteins were expressed similarly, but in minimal M9 media supplemented with 1g/L ^15^NH_4_Cl. Cells were lysed using sonication in 20 mM Tris (pH 7.5), 150 mM NaCl, 5 mM MgCl_2_, 10% (vol/vol) glycerol, 0.4% Nonidet P-40, protease inhibitors and either 1 mM DTT or 5 mM β-mercaptoethanol. Lysates were cleared by centrifugation and incubated with Ni-NTA or glutathione resin for 1-2 hours at 4 °C. After washing in high salt buffer (20 mM Tris (pH 7.5), 500 mM NaCl, 5 mM MgCl_2_ and 1 mM DTT or 5 mM β-mercaptoethanol) His-tagged proteins were eluted with 250 mM imidazole followed by thrombin cleavage. GST-fusions were cleaved with thrombin directly on the glutathione resin overnight at 4 °C. Cleaved proteins were then further purified through a S75 size exclusion column. Wild-type GTPases purified from *E. coli* are predominantly in the GDP form. Proteins were preloaded with GMPPNP or GTPγS when required. For nucleotide exchange, GTPases were incubated at 37 °C in the presence of 10 mM EDTA and a ten-fold molar excess (nucleotide:protein) of guanosine 5′-[β,γ-imido]triphosphate (GMPPNP, Sigma Aldrich), guanosine 5′-[γ-thio]triphosphate (GTPγS, Sigma Aldrich) or GTP for 10 minutes. 20 mM MgCl_2_ is then added to the sample, placed on ice, and then immediately dialyzed at 4 °C or run through an S75 size exclusion column. For hydrolysis assays, aliquots of GTP-loaded H-, K- and NRAS were flash frozen and stored at −80 °C.

### ^15^N Amino Acid Selective Labeling

Selectively labeled (^15^N-threonine, ^15^N-tyrosine, ^15^N-leucine, ^15^N-valine and ^15^N-isoleucine) proteins were expressed by culturing cells in 37 °C in M9 media supplemented with 1 g/L ^14^NH_4_Cl and 100 mg/L of the 19 unlabeled amino acids. Cultures were grown to an O.D._600_ of 0.6-0.9, at which point, 1 g/L of each of the 19 unlabeled amino acids and 100 mg/L of ^15^N-labeled amino acid. Amino acids were allowed to dissolve for 15 mins, at which point 0.25 mM IPTG was added to the culture which was then incubated overnight at 16 °C.

### NMR Spectroscopy

NMR data were recorded at 25 °C on 600 MHz Bruker UltraShield spectrometer equipped with a 5 mm CryoProbe and 1.7 mm TCI MicroCryoProbe. All multiplexed assays were performed on the 1.7 mm cryoprobe. These samples of 40 µL were prepared, with the composition and concentration of each small GTPase described in the results section and/or figure legends. Nucleotide exchange assays, GTP hydrolysis assays and GTPase/effector titrations were typically performed in 20 mM Tris (pH 7.5), 100 mM NaCl, 5-15 mM MgCl_2_ and 1 mM DTT and 10% D_2_O. Two-dimensional ^1^H/^15^N heteronuclear single quantum coherence (HSQC) spectra were collected sequentially to monitor kinetics. All multiplexed experiments used BEST-HSQCs (52). Spectra were processed with NMRPipe (53) and analyzed using NMRView (54). For intrinsic exchange and GEF assays, GTPγS was added at a 8-10 molar fold excess (GTPγS: Total Protein) and SOS_cat_ was added at molar ratios described in our results. To calculate the GDP-bound ratio [*I*_GDP_/(*I*_GDP_ + *I*_GTP_)], peak intensities were extracted from each individual spectrum using NMRView. Exchange curves were plotted and fitted to a single phase exponential decay function using GraphPad Software. For intrinsic GTP hydrolysis and GAP assays, peak intensities were extracted and data were fit to a one phase exponential association function. For effector/monobody titrations, unlabeled RBD domain or NS1 monobody was titrated into selectively labeled mixtures of GTPases.

## Acknowledgments

This work was supported by grants (to M.J.S.) from the Canadian Cancer Society Research Institute (CCSRI), Canadian Institutes for Health Research (CIHR), and the National Science and Engineering Council of Canada (NSERC). R.K. was supported by research fellowships from the Fonds de recherche du Québec – Santé (FRQS) and the Cole Foundation. M.J.S. holds a Canada Research Chair in Cancer Signalling and Structural Biology.

## Author Contributions

This project was conceived and designed by R.K. and M.J.S. Experiments were performed by R.K. The manuscript was prepared by M.J.S. and R.K.

The authors declare no conflicts of interest.

