## Supplemental Data for "Conformational resolution of nucleotide cycling and effector interactions for multiple small GTPases in parallel"

### Supplemental Figures

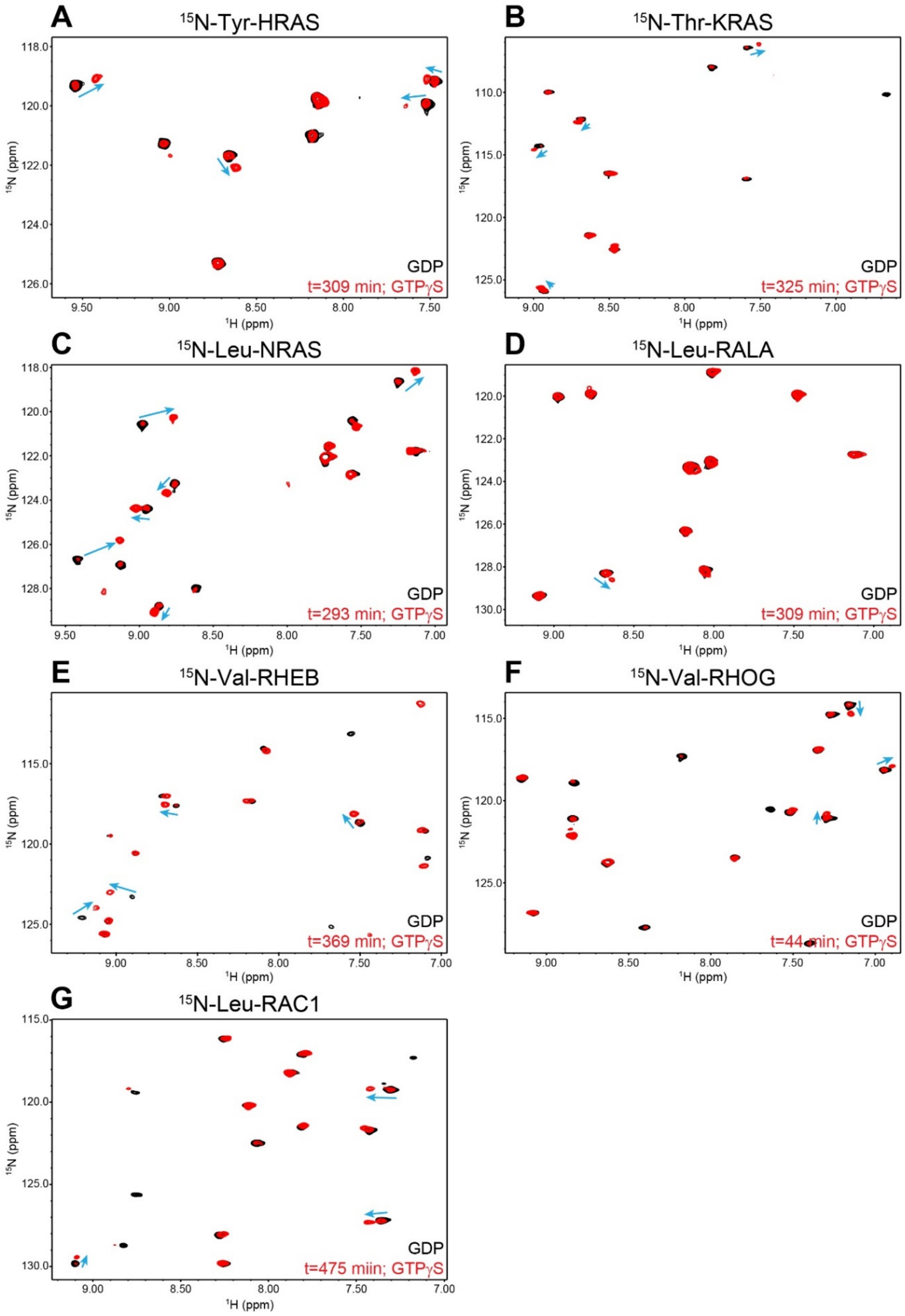

**Supplemental Fig. S1.** Representative $^1$H-$^{15}$N BEST-HSQC spectra of intrinsic nucleotide exchange assays (GDP →GTPγS) for various small GTPases, selectively labelled with specific amino acids. **A)** HRAS, Tyr labelled. **B)** KRAS, Thr labelled. **C)** NRAS, Leu labelled. **D)** RALA, Leu labelled. **E)** RHEB, Val labelled. **F)** RHOG, Val labelled. **G)** RAC1, Leu labelled. All spectra recorded at 300 μM GTPase (except RHEB, 200 μM) in 20 mM Tris, pH 7.5, 100 mM NaCl, 1 mM DTT and 10 mM $MgCl_2$ with a 10:1, GTPγS:GTPase ratio. Black spectra correspond to initial, GDP bound GTPases, and red spectra correspond to GTPγS-loaded GTPases (timepoint indicated on spectra). Arrows point to specific peaks that undergo shifts upon GDP→GTPγS exchange.

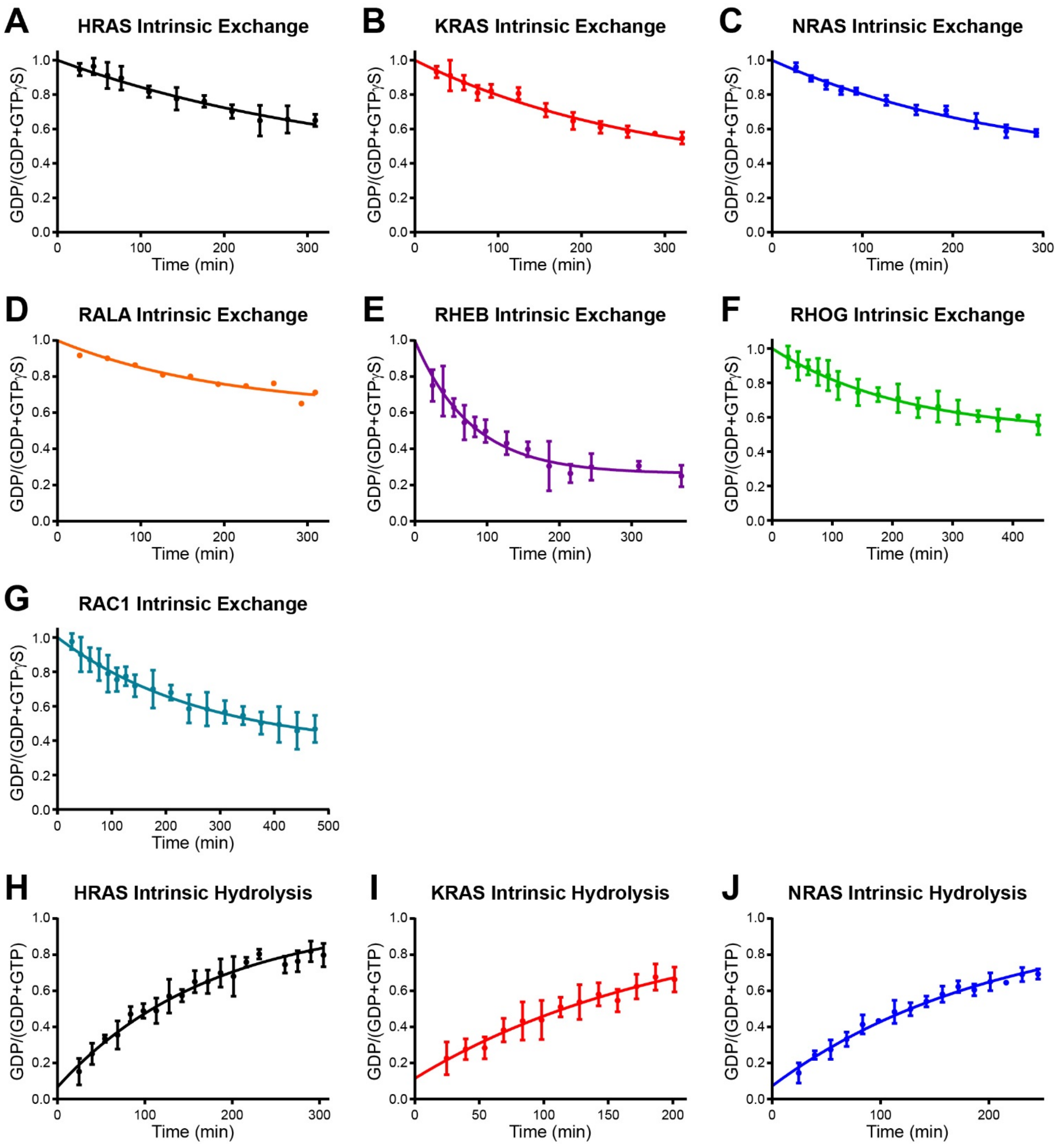


**Supplemental Fig. S2.** Nucleotide cycling assays for selectively labelled GTPases, measured individually. **A-G)** Intrinsic nucleotide exchange curves (GDP→GTPγS) for experiments conducted as described through **Supplemental Fig. S1**. **H-J)** Intrinsic GTP hydrolysis curves for the three isoforms of RAS. Experiments were performed in 20 mM Tris, pH 7.5, 100 mM NaCl, 1 mM DTT and 5 mM $MgCl_2$. The same $^{15}N$ selective labelling scheme was used as for the exchange assays.

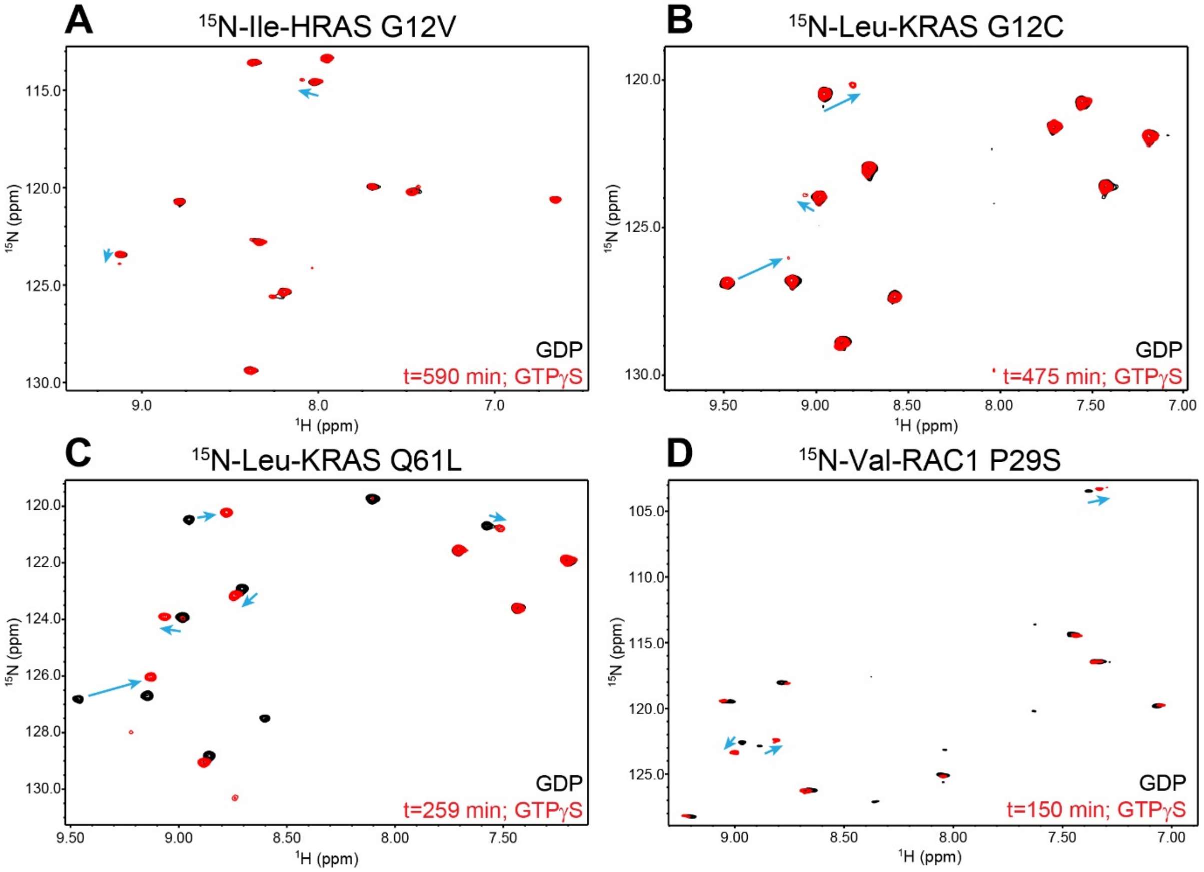


**Supplemental Fig. S3.** Representative $^{1}H$-$^{15}N$ BEST-HSQC spectra of intrinsic nucleotide exchange assays (GDP→GTPγS) for various small GTPase oncoproteins, selectively labelled with specific amino acids. **A)** HRAS G12V, Ile labelled. **B)** KRAS G12C, Leu labelled. **C)** KRAS Q61L, Leu labelled. **D)** RAC1 P29S, Val labelled. All spectra recorded at 300 μM GTPase in 20 mM Tris, pH 7.5, 100 mM NaCl, 1 mM DTT and 10 mM $MgCl_2$ with a 10:1, GTPγS:GTPase ratio. Black spectra correspond to initial, GDP bound GTPases, and red spectra correspond to GTPγS-loaded GTPases (timepoints indicated on spectra). Arrows point to specific peaks that undergo shifts upon GDP→GTPγS exchange.

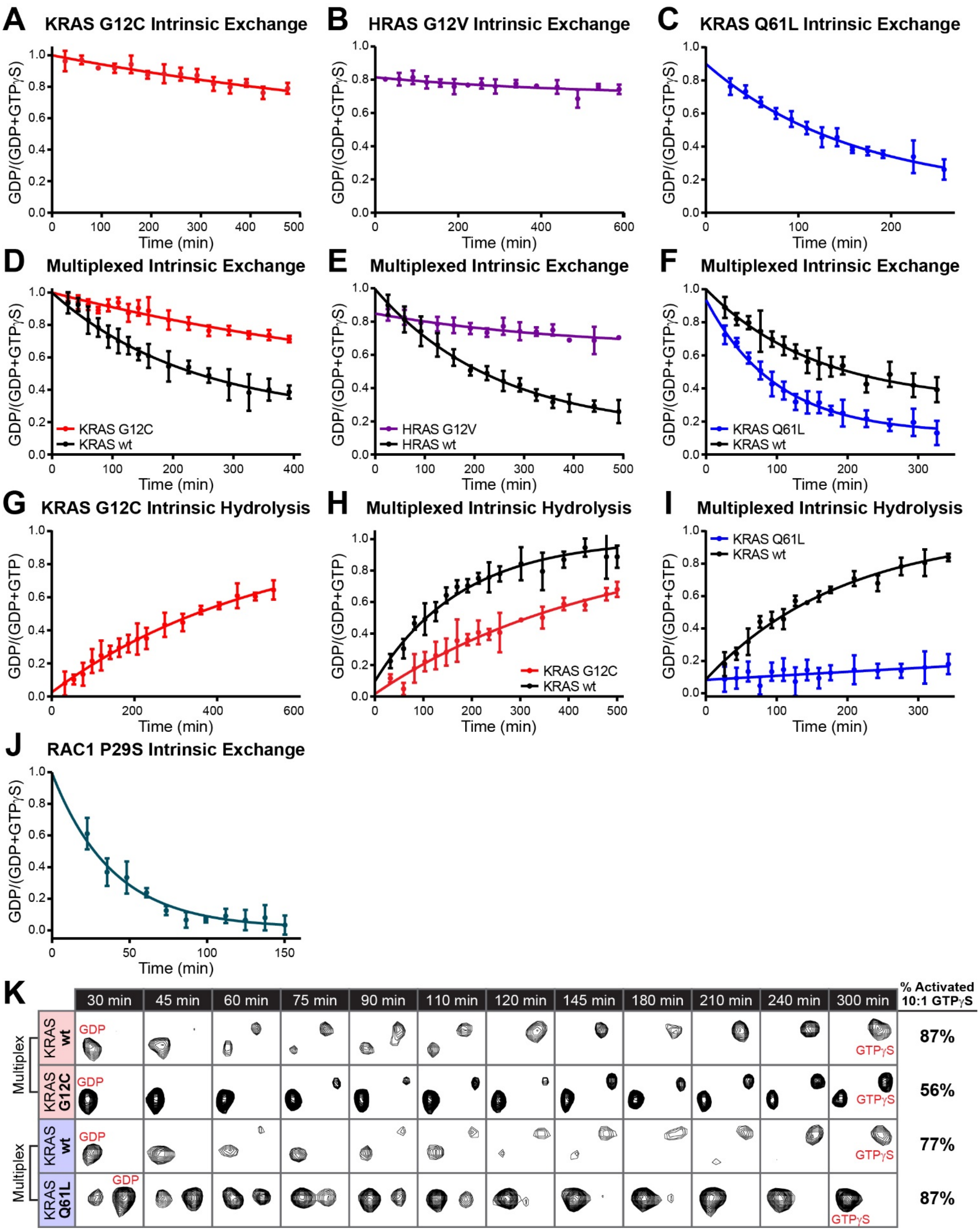


**Supplemental Fig. S4**. Nucleotide cycling assays for wild-type versus oncogenic mutant GTPases. **A)** Intrinsic nucleotide exchange curve (GDP→GTPγS) for KRAS G12C (300 μM protein, 10:1

GTPγS:GTPase, 20 mM Tris, pH 7.5, 100 mM NaCl, 1 mM DTT and 10 mM $MgCl_2$). **B)** Intrinsic nucleotide exchange curve (GDP→GTPγS) for HRAS G12V (300 µM protein, 10:1 GTPγS:GTPase, 20 mM Tris, pH 7.5, 100 mM NaCl, 1 mM DTT and 10 mM $MgCl_2$). **C)** Intrinsic nucleotide exchange curve (GDP→GTPγS) for KRAS Q61L (300 µM protein, 10:1 GTPγS:GTPase, 20 mM Tris, pH 7.5, 100 mM NaCl, 1 mM DTT and 10 mM $MgCl_2$). **D)** Multiplexed intrinsic nucleotide exchange assay with KRAS G12C and wild-type (250 µM protein each, 10:1 GTPγS:GTPase, 20 mM Tris, pH 7.5, 100 mM NaCl, 1 mM DTT and 10 mM $MgCl_2$). **E)** Multiplexed intrinsic nucleotide exchange assay with HRAS G12V and wild-type (300 µM protein each, 10:1 GTPγS:GTPase, 20 mM Tris, pH 7.5, 100 mM NaCl, 1 mM DTT and 10 mM $MgCl_2$). **F)** Multiplexed intrinsic nucleotide exchange assay with KRAS Q61L and wild-type (250 µM protein each, 10:1 GTPγS:GTPase, 20 mM Tris, pH 7.5, 100 mM NaCl, 1 mM DTT and 10 mM $MgCl_2$). **G)** Intrinsic GTP hydrolysis curve of KRAS G12C (150 µM protein, 20 mM Tris, pH 7.5, 100 mM NaCl, 1 mM DTT and 5 mM $MgCl_2$). **H)** Multiplexed intrinsic GTP hydrolysis assay with KRAS G12C and wild-type (125 µM protein each, 20 mM Tris, pH 7.5, 100 mM NaCl, 1 mM DTT and 5 mM $MgCl_2$). **I)** Multiplexed intrinsic GTP hydrolysis assay with KRAS Q61L and wild-type (125 µM protein each, 20 mM Tris, pH 7.5, 100 mM NaCl, 1 mM DTT and 5 mM $MgCl_2$). **J)** Intrinsic nucleotide exchange curve (GDP →GTPγS) for RAC1 P29S (300 µM protein, 10:1 GTPγS:GTPase, 20 mM Tris, pH 7.5, 100 mM NaCl, 1 mM DTT and 10 mM $MgCl_2$). **K)** Snapshots of select resonances from KRAS wild-type/G12C multiplexed assay and KRAS wild-type/Q61L multiplexed assay. These peaks are from multiplexed assays looking at intrinsic nucleotide exchange with these oncoproteins and their wild-type counterpart.

**Supplemental Table S1.** NMR-derived hydrolysis and exchange rates for GTPases measured individually. Rates are recorded as min$^{-1}$. '*% Activated*' corresponds to the percentage of GTPase loaded with GTPγS at equilibrium, derived from the plateau in a one-phase decay fit. '*GDP Preference*' is the fold bias towards binding GDP over GTPγS, calculated using the equation: (% GDP-bound / % GTPγS-bound)*([GTPγS] / [GDP]).

| | GTPase | Reaction | Concentration | Mg$^{2+}$ | Isotopic Label | Nucleotide | Rate (x 10$^{-3}$min$^{-1}$) | % Activated | GDP Preference (fold$^{GTP\gamma S}$) |
|---|---|---|---|---|---|---|---|---|---|
| | **Supplemental Fig. S2** | | | | | | | | |
| A | HRAS | Exchange | 300 µM | 10 mM | $^{15}$N-Tyr | 10:1 GTPγS | 2.7 ± 0.9 | 66 ± 17 | 5.2x |
| B | KRAS | Exchange | 300 µM | 10 mM | $^{15}$N-Thr | 10:1 GTPγS | 3.5 ± 0.7 | 68 ± 9 | 4.7x |
| C | NRAS | Exchange | 300 µM | 10 mM | $^{15}$N-Leu | 10:1 GTPγS | 3.7 ± 0.7 | 63 ± 8 | 5.9x |
| D | RALA | Exchange | 300 µM | 10 mM | $^{15}$N-Leu | 10:1 GTPγS | 5.0 ± 1.7 | 38 ± 7 | 16.3x |
| E | RHEB | Exchange | 200 µM | 10 mM | $^{15}$N-Val | 10:1 GTPγS | 12.8 ± 0.9 | 74 ± 2 | 3.5x |
| F | RHOG | Exchange | 300 µM | 10 mM | $^{15}$N-Val | 10:1 GTPγS | 4.6 ± 0.4 | 49 ± 2 | 10.4x |
| G | RAC1 | Exchange | 300 µM | 10 mM | $^{15}$N-Leu | 10:1 GTPγS | 3.7 ± 0.3 | 65 ± 3 | 5.4x |
| H | HRAS | Hydrolysis | 200 µM | 5 mM | $^{15}$N-Tyr | - | 5.7 ± 0.2 | - | - |
| I | KRAS | Hydrolysis | 200 µM | 5 mM | $^{15}$N-Thr | - | 5.0 ± 0.2 | - | - |
| J | NRAS | Hydrolysis | 200 µM | 5 mM | $^{15}$N-Leu | - | 4.8 ± 0.2 | - | - |
| | **Fig. 2** | | | | | | | | |
| A | HRAS | Exchange | 150 µM | 5 mM | $^{15}$N-uniform | 10:1 GTPγS | 2.3 ± 0.5 | 59 ± 9 | 6.9x |
| | | Exchange | 250 µM | 5 mM | $^{15}$N-uniform | 10:1 GTPγS | 2.6 ± 0.5 | 72 ± 10 | 3.9x |
| | | Exchange | 350 µM | 5 mM | $^{15}$N-uniform | 10:1 GTPγS | 4.1 ± 0.2 | 81 ± 2 | 2.3x |
| B | KRAS | Exchange | 150 µM | 5 mM | $^{15}$N-uniform | 10:1 GTPγS | 4.2 ± 0.5 | 60 ± 4 | 6.7x |
| | | Exchange | 250 µM | 5 mM | $^{15}$N-uniform | 10:1 GTPγS | 4.2 ± 0.3 | 71 ± 3 | 4.1x |
| | | Exchange | 350 µM | 5 mM | $^{15}$N-uniform | 10:1 GTPγS | 7.2 ± 0.2 | 82 ± 1 | 2.2x |
| C | NRAS | Exchange | 150 µM | 5 mM | $^{15}$N-uniform | 10:1 GTPγS | 2.7 ± 0.3 | 63 ± 5 | 5.9x |
| | | Exchange | 250 µM | 5 mM | $^{15}$N-uniform | 10:1 GTPγS | 3.5 ± 0.3 | 70 ± 4 | 4.3x |
| | | Exchange | 350 µM | 5 mM | $^{15}$N-uniform | 10:1 GTPγS | 6.5 ± 0.2 | 76 ± 1 | 3.2x |
| D | NRAS | Exchange | 200 µM | 5 mM | $^{15}$N-uniform | 10:1 GTPγS | 2.5 ± 0.1 | 63 ± 18 | 5.9x |
| | | Exchange | 350 µM | 5 mM | $^{15}$N-uniform | 10:1 GTPγS | 5.9 ± 0.3 | 82 ± 2 | 2.2x |
| | | Exchange | 350 µM | 15 mM | $^{15}$N-uniform | 10:1 GTPγS | 2.7 ± 0.9 | 69 ± 17 | 4.5x |
| F | KRAS G13D | Exchange | 250 µM | 5 mM | $^{15}$N-uniform | 10:1 GTPγS | 31.3 ± 3.7 | 86 ± 3 | 1.6x |
| | | Exchange | 250 µM | 50 mM | $^{15}$N-uniform | 10:1 GTPγS | 28.5 ± 3.2 | 85 ± 3 | 1.8x |
| F | KRAS Q61L | Exchange | 300 µM | 5 mM | $^{15}$N-Leu | 10:1 GTPγS | 9.2 ± 0.9 | 90 ± 3 | 1.1x |
| | | Exchange | 300 µM | 50 mM | $^{15}$N-Leu | 10:1 GTPγS | 8.5 ± 0.8 | 89 ± 3 | 1.2x |
| G | HRAS | Hydrolysis | 120 µM | 5 mM | $^{15}$N-uniform | - | 10 ± 0.6 | - | - |
| | | Hydrolysis | 220 µM | 5 mM | $^{15}$N-uniform | - | 8.7 ± 0.3 | - | - |
| H | KRAS | Hydrolysis | 150 µM | 5 mM | $^{15}$N-uniform | - | 8.7 ± 0.5 | - | - |
| | | Hydrolysis | 250 µM | 5 mM | $^{15}$N-uniform | - | 9.0 ± 0.4 | - | - |
| I | NRAS | Hydrolysis | 115 µM | 5 mM | $^{15}$N-uniform | - | 7.4 ± 0.3 | - | - |
| | | Hydrolysis | 190 µM | 5 mM | $^{15}$N-uniform | - | 8.3 ± 0.2 | - | - |
| | **Supplemental Fig. S4** | | | | | | | | |
| A | KRAS G12C | Exchange | 300 µM | 10 mM | $^{15}$N-Leu | 10:1 GTPγS | 1.1 ± 0.9 | 55 ± 38 | 8.2x |
| B | HRAS G12V | Exchange | 300 µM | 10 mM | $^{15}$N-Ile | 10:1 GTPγS | 3.6 ± 0.3 | 27 ± 3 | 27.0x |
| C | KRAS Q61L | Exchange | 300 µM | 10 mM | $^{15}$N-Leu | 10:1 GTPγS | 8.6 ± 0.1 | 81 ± 4 | 2.3x |
| D | KRAS G12C | Hydrolysis | 150 µM | 5 mM | $^{15}$N-Leu | - | 1.9 ± 0.1 | - | - |
| E | RAC1 P29S | Exchange | 300 µM | 10 mM | $^{15}$N-Val | 10:1 GTPγS | 25.6 ± 2.1 | 98 ± 2 | 0.2x |

**Supplemental Table S2.** NMR-derived rates for multiplexed experiments. Rates are recorded as min$^{-1}$. '*% Activated*' corresponds to % of GTPase loaded with GTPγS at equilibrium, derived from the plateau in a one-phase decay fit. '*GDP Preference*' is the fold bias towards binding GDP over GTPγS, calculated using the equation: (% GDP-bound / % GTPγS-bound)*([GTPγS] / [GDP]).

| | GTPase | Reaction | Concentration | Mg$^{2+}$ | Isotopic Label | GAP/ GEF | Ratio | Nucleotide | Rate (x 10$^{-3}$ min$^{-1}$) | % Activated | GDP Preference (fold$^{GTP\gamma S}$) |
|---|---|---|---|---|---|---|---|---|---|---|---|
| **Fig. 3** | | | | | | | | | | | |
| E | HRAS | Exchange | 300 µM | 15 mM | $^{15}$N-Tyr | - | - | 8:1 GTPγS | 3.5 ± 0.3 | 80 ± 4 | 2.0x |
| | KRAS | | | | $^{15}$N-Thr | | | | 5.7 ± 0.5 | 81 ± 3 | 1.9x |
| | NRAS | | | | $^{15}$N-Leu | | | | 3.9 ± 0.3 | 90 ± 3 | 0.9x |
| J | NRAS | Exchange | 300 µM | 15 mM | $^{15}$N-Leu | - | - | 8:1 GTPγS | 3.9 ± 0.6 | 84 ± 6 | 1.5x |
| | RALA | | | | $^{15}$N-Leu | | | | 5.3 ± 1.6 | 46 ± 4 | 9.4x |
| | RHOG | | | | $^{15}$N-Val | | | | 5.1 ± 0.5 | 49 ± 2 | 8.3x |
| **Fig. 4** | | | | | | | | | | | |
| B | HRAS | Exchange | 300 µM | 15 mM | $^{15}$N-Tyr | SOScat | 1:15000 | 8:1 GTPγS | 15.7 ± 1.6 | 77 ± 3 | 2.4x |
| | KRAS | | | | $^{15}$N-Thr | | | | 18.6 ± 1.9 | 80 ± 3 | 2.0x |
| | NRAS | | | | $^{15}$N-Leu | | | | 11.9 ± 1.2 | 94 ± 4 | 0.5x |
| C | NRAS | Exchange | 300 µM | 15 mM | $^{15}$N-Leu | SOScat | 1:30000 | 8:1 GTPγS | 6.1 ± 0.5 | 96 ± 3 | 0.3x |
| | RALA | | | | $^{15}$N-Leu | | | | 7.7 ± 4.9 | 39 ± 5 | 12.5x |
| | RHOG | | | | $^{15}$N-Val | | | | 5.1 ± 1.0 | 45 ± 4 | 9.8x |
| D | KRAS | Exchange | 300 µM | 10 mM | $^{15}$N-Thr | - | - | 10:1 GTPγS | 7.0 ± 0.7 | 83 ± 4 | 2.0x |
| | RAC1 | | | | $^{15}$N-Leu | | | | 5.5 ± 0.4 | 65 ± 3 | 5.4x |
| E | KRAS | Exchange | 300 µM | 10 mM | $^{15}$N-Thr | SOScat | 1:8000 | 10:1 GTPγS | 16.9 ± 1.2 | 83 ± 2 | 2.0x |
| | RAC1 | | | | $^{15}$N-Leu | | | | 6.5 ± 0.5 | 67 ± 3 | 4.9x |
| F | HRAS | Hydrolysis | 125 µM | 5 mM | $^{15}$N-Tyr | - | - | - | 4.3 ± 0.3 | - | - |
| | KRAS | | | | $^{15}$N-Thr | | | | 4.5 ± 0.3 | - | - |
| | NRAS | | | | $^{15}$N-Leu | | | | 4.5 ± 0.1 | - | - |
| H | HRAS | Hydrolysis | 125 µM | 5 mM | $^{15}$N-Tyr | GAP-334 | 1:5500 | - | 10.0 ± 1.1 | - | - |
| | KRAS | | | | $^{15}$N-Thr | | | | 11.6 ± 1.1 | - | - |
| | NRAS | | | | $^{15}$N-Leu | | | | 12.9 ± 0.8 | - | - |
| **Supplemental Fig. S4** | | | | | | | | | | | |
| D | KRAS WT | Exchange | 250 µM | 10 mM | $^{15}$N-Thr | - | - | 10:1 GTPγS | 4.4 ± 0.4 | 77 ± 4 | 3.0x |
| | KRAS G12C | | | | $^{15}$N-Leu | | | | 1.5 ± 0.8 | 67 ± 29 | 4.9x |
| E | HRAS WT | Exchange | 300 µM | 10 mM | $^{15}$N-Tyr | - | - | 10:1 GTPγS | 4.1 ± 0.2 | 86 ± 2 | 1.6x |
| | HRAS G12V | | | | $^{15}$N-Ile | | | | 2.5 ± 1.8 | 36 ± 9 | 17.8x |
| F | KRAS WT | Exchange | 250 µM | 10 mM | $^{15}$N-Thr | - | - | 10:1 GTPγS | 6.4 ± 0.6 | 69 ± 3 | 4.5x |
| | KRAS Q61L | | | | $^{15}$N-Leu | | | | 10.1 ± 0.9 | 87 ± 2 | 1.5x |
| H | KRAS WT | Hydrolysis | 125 µM | 5 mM | $^{15}$N-Thr | - | - | - | 5.6 ± 0.3 | - | - |
| | KRAS G12C | | | | $^{15}$N-Leu | | | | 2.1 ± 0.1 | - | - |
| I | KRAS WT | Hydrolysis | 125 µM | 5 mM | $^{15}$N-Thr | - | - | - | 5.2 ± 0.2 | - | - |
| | KRAS Q61L | | | | $^{15}$N-Leu | | | | Not observed | - | - |
| **Fig. 5** | | | | | | | | | | | |
| B | KRAS WT | Exchange | 250 µM | 10 mM | $^{15}$N-Thr | SOScat | 1:8000 | 10:1 GTPγS | 12.3 ± 1.1 | 87 ± 3 | 1.5x |
| | KRAS G12C | | | | $^{15}$N-Leu | | | | 8.1 ± 0.7 | 56 ± 2 | 7.9x |
| C | KRAS WT | Exchange | 250 µM | 5 mM | $^{15}$N-Thr | SOScat | 1:8000 | 10:1 GTPγS | 10.9 ± 1.1 | 77 ± 3 | 3.0x |
| | KRAS Q61L | | | | $^{15}$N-Leu | | | | 14.5 ± 1.4 | 87 ± 1 | 1.5x |
| D | KRAS WT | Hydrolysis | 125 µM | 5 mM | $^{15}$N-Thr | GAP-334 | 1:5000 | - | 13.4 ± 1.2 | - | - |
| | KRAS G12C | | | | $^{15}$N-Leu | | | | 2.1 ± 0.1 | - | - |
| E | KRAS WT | Hydrolysis | 125 µM | 5 mM | $^{15}$N-Thr | GAP-334 | 1:5000 | - | 13.9 ± 0.9 | - | - |
| | KRAS Q61L | | | | $^{15}$N-Leu | | | | Not observed | - | - |
| G | RAC1 WT | Exchange | 250 µM | 10 mM | $^{15}$N-Leu | - | - | 10:1 GTPγS | 3.1 ± 1.1 | 58 ± 15 | 7.2x |
| | RAC1 P29S | | | | $^{15}$N-Val | | | | 26.7 ± 1.8 | 97 ± 1 | 0.3x |

**Supplemental Table S3.** Compilation of nucleotide and $Mg^{2+}$ binding affinity to various small GTPases, as gathered from previous literature.

| | | | | Equilibrium Binding Affinity ($K_d$) | | | | | |
|---|---|---|---|---|---|---|---|---|---|
| **Reference** | **GTPase** | **Variant** | **Technique** | **GDP** | **GTP** | **GTPγS** | **GMPPNP** | **$Mg^{2+}$** | **[$Mg^{2+}$]** |
| **Sigal *et al*, 1986** (1) | HRAS | wild-type | filter binding | 20 nM | 10 nM | - | - | - | 1.5 mM |
| | HRAS | G12V | filter binding | 20 nM | 10 nM | - | - | - | |
| **Tucker *et al*, 1986** (2) | HRAS | wild-type | unknown | - | $1.5X^{GDP}$ | $0.29X^{GDP}$ | - | - | unknown |
| **Feuerstein *et al*, 1987** (3) | HRAS | wild-type | filter binding | 17.5 pM | - | - | - | - | 5 mM |
| | HRAS | G12V | | 14.1 pM | - | - | - | - | |
| **Trahey *et al*, 1987** (4) | NRAS | wild-type | filter binding | 13 nM | 12 nM | - | - | - | 5 mM |
| | NRAS | G12D | | 9 nM | 8 nM | - | - | - | |
| | NRAS | G12V | | 6 nM | 9 nM | - | - | - | |
| **Feig *et al*, 1988** (5) | HRAS | wild-type | filter binding | - | 8 nM | - | - | - | 1 mM |
| | HRAS | S17N | | 5 nM | 300 nM | - | - | - | |
| **Feuerstein *et al*, 1989** (6) | HRAS | wild-type | filter binding | - | 10.6 pM | 34.5 pM | - | - | 10 mM |
| **John *et al*, 1989** (7) | HRAS | wild-type | filter binding | - | $1.9X^{GDP}$ | $0.72X^{GDP}$ | $0.09X^{GDP}$ | - | 10 mM |
| | HRAS | G12V | | - | $0.67X^{GDP}$ | $0.37X^{GDP}$ | $0.035X^{GDP}$ | - | |
| **Scherer *et al*, 1989** (8) | HRAS | wild-type | filter binding | - | $1.0X^{GDP}$ | $0.7X^{GDP}$ | $0.09X^{GDP}$ | - | 10 mM |
| **John *et al*, 1990** (9) | HRAS | wild-type | filter binding | 1.6 pM | 0.56 pM | - | - | - | 5 mM |
| **Kabcenell *et al*, 1990** (10) | ySec4p | wild-type | filter binding | 77 nM | 3.5 nM | 3.7 nM | - | - | 5 mM |
| **Reinstein *et al*, 1991** (11) | HRAS | wild-type | filter binding/ mantGDP | 1.6 pM | - | - | - | - | 10 mM |
| | HRAS | F28L | | 182 pM | - | - | - | $15.2\ \mu M^{GDP}$<br>$0.03\ \mu M^{GTP}$<br>$0.02\ \mu M^{GMPPNP}$ | |
| **Burstein *et al*, 1992** (12) | RAB3A | wild-type | filter binding | 63 nM | - | 46 nM | - | $4\ \mu M^{free}$ | 10 mM |
| **John *et al*, 1993** (13) | HRAS | wild-type | filter/mantGDP | 20.3 pM | 10.7 pM | - | - | $2.8\ \mu M^{GDP}$ | 10 mM |
| **Rensland *et al*, 1995** (14) | HRAS | wild-type | filter binding | - | $1.9X^{GDP}$ | - | $0.09X^{GDP}$ | - | unknown |
| **Ahmadian *et al*, 1999** (15) | HRAS | wild-type | unknown | - | - | - | 0.24 nM | - | unknown |
| **Zhang *et al*, 2000** (16) | RAC1 | wild-type | filter binding | 620 nM | - | 240 nM | - | - | 10 nM |
| | CDC42 | wild-type | | 590 nM | - | 170 nM | - | - | |
| | RHOA | wild-type | | 480 nM | - | 160 nM | - | - | |
| **Ford *et al*, 2009** (17) | HRAS | wild-type | mantGDP/GTP | 68.0 pM | 79.1 pM | - | - | - | 5 mM |
| | HRAS | G60A | | 22.3 pM | 36.1 pM | - | - | - | |
